# A Deformulating Multi-Attribute SEC Method for Simultaneous Quantification of Total RNA, Conjugated Antibody, and Free Antibody in Targeted Lipid Nanoparticles

**DOI:** 10.64898/2026.11.06.764440

**Authors:** Jeremy R. Filteau, Anastasiya P. Manuilov, Garret M. Rubin, Hui Zhou, Paroma Ghosh, Mohui Wei, Liang Xu, Himanshu Dhamankar, Peter Bernhardt

## Abstract

Targeted lipid nanoparticles (tLNPs) hold promise for improving cell- and tissue-selective delivery of therapeutic payloads but introduce analytical challenges associated with characterizing both encapsulated cargo and surface-associated targeting ligands within the same formulation. Conventional nucleic acid and protein assays can be affected by nanoparticle scattering, matrix interference, or a limited ability to distinguish conjugated antibody from free antibody. Here, we report an online deformulating size-exclusion high-performance liquid chromatography (SE-HPLC) method that enables simultaneous quantification of total RNA, conjugated antibody, and free antibody from 5 μL of tLNP sample. Following injection, exposure to a mobile phase containing sodium dodecyl sulfate and isopropanol disrupts the nanoparticles, enabling chromatographic separation of RNA, conjugated antibody, and free (unconjugated) antibody while minimizing sample handling prior to analysis. RNA was quantified by UV260 detection, whereas antibody was quantified by UV280 or intrinsic fluorescence detection to maximize sensitivity. The method demonstrated linear response over the evaluated concentration ranges, spike recoveries of 88-122%, and precision below 5% relative standard deviation. Application to purified tLNP formulations demonstrated simultaneous assessment of total RNA and conjugated antibody content, while confirming the absence of detectable free antibody following purification. Together, these results establish online deformulating SE-HPLC as a practical multi-attribute analytical method for characterization of tLNPs, enabling concurrent assessment of total RNA concentration, conjugated antibody concentration, and free antibody concentration using standard HPLC instrumentation.

## Intro/Background

Lipid nanoparticles (LNPs) have emerged as a leading nonviral platform for nucleic acid delivery, combining protection from degradation with efficient intracellular delivery of therapeutic cargo.^1–3^ The clinical success of LNP-formulated RNA vaccines for COVID19 demonstrated the safety, efficacy and manufacturability of the platform on a global scale.^4,5^ Current applications include protein replacement, gene editing, RNA interference, cancer immunotherapy, and cell- or tissue-specific delivery.^1,2,6^ As applications expand, increasingly sophisticated LNP formulations are being developed to modulate biodistribution and enhance delivery to specific biological targets. Systemically administered LNPs often exhibit substantial liver tropism, which can limit extrahepatic and cell-selective delivery.^7^ One strategy for redirecting biodistribution is to modify the chemical identity or relative abundance of formulation lipids.^8–10^ A complementary strategy is to functionalize the nanoparticle surface with ligands that recognize receptors or antigens on cells of interest.^11–13^ Antibodies are attractive targeting ligands because their affinity and specificity can be engineered to direct LNP cargo to selected tissues and cell populations.^11,14–17^ Such targeted LNPs (tLNPs) are commonly prepared by coupling proteins or peptides to lipids bearing reactive groups such as maleimide or dibenzocyclooctyne (DBCO), either before or after nanoparticle formation. The addition of a surface-bound targeting ligand introduces attributes related to ligand abundance, conjugation efficiency, and payload-to-ligand ratios. Consequently, the development of tLNPs requires analytical methods capable of accurately quantifying total RNA, conjugated antibody, and free antibody within the same formulated sample.

Although assays exist for nucleic acid and protein characterization, most were developed for measurement of individual analytes and may be susceptible to matrix effects when applied to complex tLNP formulations. While direct UV measurement is routinely used for isolated nucleic acids and proteins, application to nanoparticle formulations is complicated by particle-associated light scattering and other matrix interferences that disguise the true analyte concentrations. The RiboGreen assay is commonly used to quantify total nucleic acid in LNPs through detergent-mediated payload release and binding of a fluorescent dye. However, assay response can depend on formulation composition and sample-preparation conditions.^18,19^ Bradford and bicinchoninic acid assays for protein quantification may be affected by formulation additives, lipids, and RNA, limiting their use in direct measurement of tLNPs.^20,21^ These assays are often used to estimate surface-associated antibody indirectly through measurement of residual antibody in the reaction supernatant, but this is not ideal. Sodium dodecyl sulfate-polyacrylamide gel electrophoresis can directly confirm protein conjugation and is generally less susceptible to interference than colorimetric protein assays, but it is difficult to provide accurate quantification.^22^ Immunoassays offer molecular specificity and reduced matrix interference, but may underestimate surface-displayed protein if steric constraints limit access to the relevant epitope.^23^ None of these approaches can distinguish and quantify total RNA, conjugated antibody, and free antibody within a single measurement or assay run. As a result, tLNP characterization frequently requires assembling information from several independent assays to obtain a complete picture of product composition, increasing sample consumption and analytical workload. An ideal method would minimize sample handling and matrix interference, while simultaneously quantifying total RNA, conjugated antibody, and free antibody within a single analytical workflow.

Recently, a deformulating size-exclusion high-performance liquid chromatography (SE-HPLC) method was developed for unconjugated LNPs, enabling online nanoparticle deformulation and quantification of nucleic acid payloads.^24^ This work demonstrated a streamlined approach for analysis of nucleic acid-loaded LNPs while reducing the influence of formulation-related interferences. Given the widespread use of SE-HPLC for protein characterization, extending this approach to tLNPs represents a promising strategy to achieve simultaneous analysis of nucleic acid and antibody species.^25–27^ In addition to quantifying total antibody, differentiation of conjugated and free antibody is desirable because the relative abundance of these species may influence product quality and biological performance. To our knowledge, no analytical method has enabled the simultaneous quantification of total RNA, conjugated antibody, and free antibody from a single measurement.

Here, we describe a deformulating SE-HPLC method for the simultaneous quantification of total RNA, conjugated antibody, and free antibody in antibody-functionalized LNP samples. Exposure to an SDS/IPA containing mobile phase disrupts the nanoparticles online, releasing RNA and antibody species for chromatographic separation. Multiwavelength UV detection enabled independent quantification of RNA and antibody at their respective absorbance maxima (260 and 280 nm), while intrinsic fluorescence detection provides enhanced sensitivity for antibody measurement. The retention time difference between free and conjugated antibody made quantification of both species possible under the tested conditions. Following establishment of the chromatographic method, specificity, linearity, precision, and accuracy were evaluated across representative concentration ranges. This approach extends the utility of SE-HPLC beyond the separate characterization of nucleic acid-loaded LNPs and protein therapeutics, enabling the simultaneous characterization of both nucleic acid and antibody species within a single targeted LNP (tLNP) formulation

## Materials/Methods

Chemicals and solvents used include 10x phosphate buffered saline (PBS) from Fisher Scientific containing no calcium or magnesium, isopropanol (IPA) Optima LC-MS grade from Fisher Chemical, UHPLC grade water from Fisher Chemical, and sodium dodecyl sulfate (SDS) from Sigma-Aldrich. LNPs shown in the manuscript were prepared under proprietary conditions. The identities of the RNA and antibody used in the manuscript are confidential.

### Deformulating SE-HPLC Conditions

Experiments were conducted using a Waters ACQUITY UPLC with UV detection (260 nm or 280 nm) and fluorescence detection (280 nm excitation, 350 nm emission). A Waters GTxResolve Premier BEH SEC 450Å, 2.5 μm, 4.6 × 300 mm column was used for online deformulation and separation. The optimized mobile phase was 1x phosphate buffered saline (10 mM phosphate, 137 mM NaCl, 2.7 mM KCl, pH 7.4) 20% isopropanol, and 0.4% SDS (w/v), delivered isocratically at 0.25 mL/min with column temperature of 45 °C. The injection volume was 5 μL with a runtime of 20 min per injection, including equilibration. Injections were performed in triplicate and the average was reported unless otherwise specified.

### Preparation of Calibration Standards

RNA, free (unconjugated) antibody, and empty LNPs were combined to prepare calibration standards used in linearity assessment and subsequent quantification of tLNPs. A constant amount of empty LNP was included in each standard level to approximate the lipid matrix present in tLNP samples. The dilution buffer used was 10 mM PBS. UV calibration levels ranged from 5 to 200 μg/mL for both analytes; fluorescence antibody calibration levels ranged from 1.25 to 200 μg/mL. Samples were transferred to glass vials and stored at 8 °C in the autosampler until analysis.

### Preparation of Spike-Recovery Samples

Known amounts of RNA and free antibody were spiked into a representative tLNP sample to generate spike amounts between 1.25–50.0 μg/mL. The dilution buffer used was 10 mM PBS. A fraction of representative tLNP was left unspiked to establish baseline concentration of RNA and antibody. Samples were transferred to glass vials and stored at 8 °C in the autosampler until analysis.

### Data Analysis

Chromatograms were processed in Empower CDS. Peak assignments were established using individual reference standards. Analyte concentrations were determined from external calibration using linear regression, and calibration curves were not forced through the origin. Retention time and peak area precision were assessed from triplicate injections. Perpendicular-drop integration was applied when necessary to resolve overlapping peaks. Back-calculated recovery (%) was calculated as the concentration predicted from the calibration curve divided by the nominal concentration and multiplied by 100. Percent recovery (%) was calculated as the measured concentration divided by the expected concentration and multiplied by 100. Relative standard deviation (RSD) was calculated from replicate measurements using the mean and standard deviation. RNA-to-antibody ratio was calculated on a mass basis by dividing the measured RNA concentration by the measured antibody concentration.

## Results/Discussion

### Method Development and Optimization

Intact tLNPs were injected directly into the SE-HPLC system, where exposure to the SDS/IPA containing mobile phase deformulates the nanoparticles, releasing RNA and antibody species for chromatographic separation and quantification (**Figure 1A**). The tLNPs in this work utilized a site-specific antibody conjugation to enable controlled display on the nanoparticle surface. Under the chromatographic conditions employed, low-molecular-weight formulation components, including residual lipids, excipients, salts, and buffer constituents, elute near the total permeation volume of the column, avoiding interference with RNA or antibody quantification. RNA was quantified by UV260, while antibody was quantified by UV280 or intrinsic fluorescence (Ex 280 nm, Em 350 nm). Several SE-HPLC columns were evaluated under deformulating mobile phase conditions of 1× phosphate-buffered saline with 0.4% SDS/20% IPA at a column temperature of 45°C. Of the tested columns, the Waters GTxResolve Premier 450Å, 4.6 × 300 mm provided the best balance of sensitivity, runtime, and resolution and was selected for future use (**Supplemental Figure S1A–B**).

**Figure 1.**
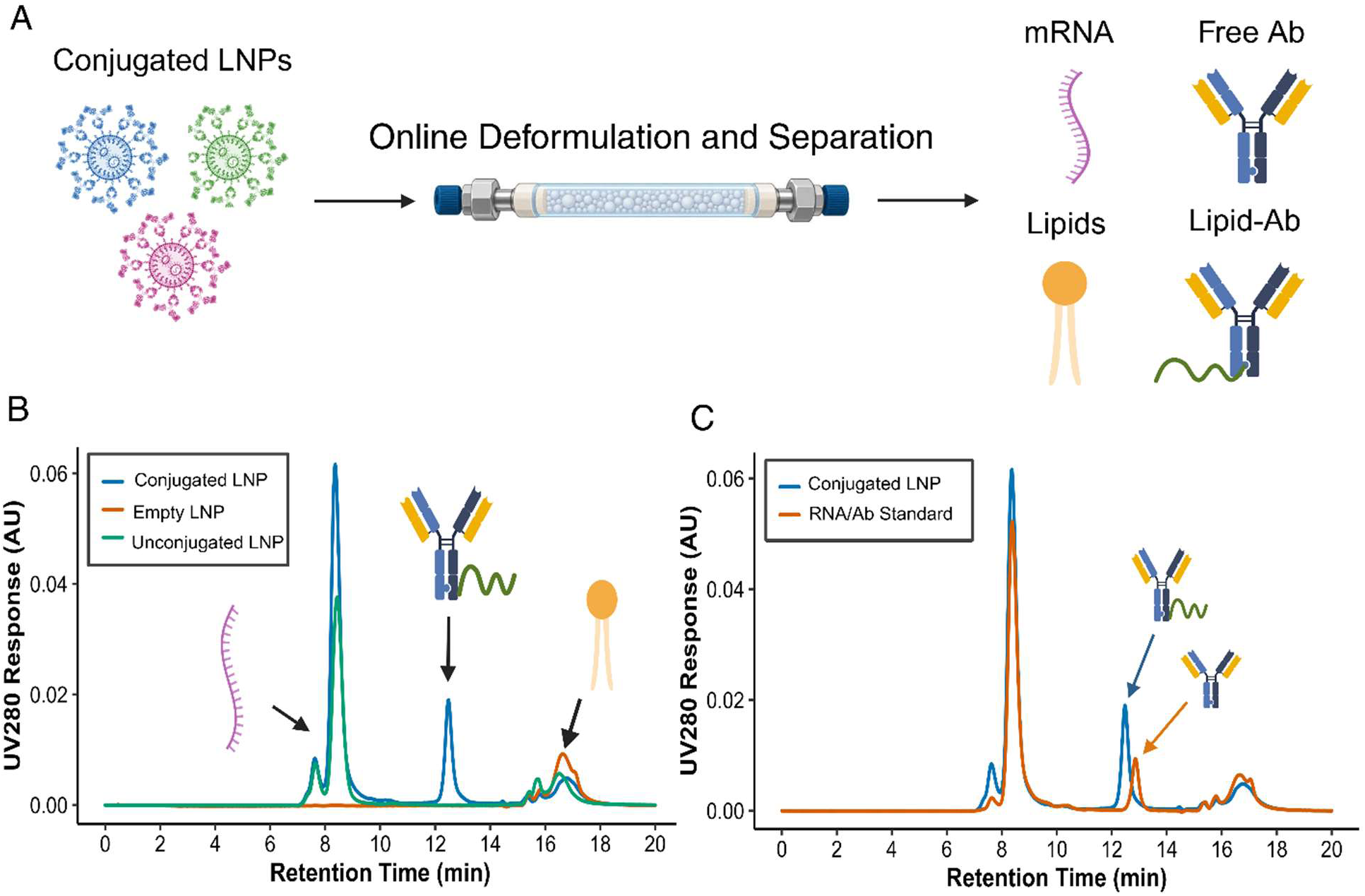
Online deformulation SE-HPLC workflow and specificity of separated components. (A) Schematic of direct tLNP injection, deformulation, and SE-HPLC separation of tLNP components. (B) UV280 chromatograms of conjugated LNP, Unconjugated LNP, and empty LNP controls demonstrating specificity of RNA and antibody signal regions. (C) Overlay of a tLNP chromatogram with an RNA/free-antibody reference standard. Arrows indicate the retention-time difference between conjugated and free antibody.

To test the extent of deformulation under these conditions, a tLNP sample was manually spiked to 0.4% SDS/20% IPA, vortexed for 30 seconds at max intensity, and promptly measured alongside an untreated sample by SE-HPLC. We observed comparable RNA peak areas between manually deformulated samples and tLNPs deformulated online under the method conditions, indicating that additional sample disruption did not improve RNA recovery (**Supplemental Figure S1C**). This aligns with previous SE-HPLC results, where 0.2% SDS/20% isopropanol was found to be highly effective for online deformulation of several tested LNPs.^24^ In the present work, 0.4% SDS was employed to provide an additional margin for complete deformulation and analyte solubilization. Unlike conventional nucleic acid LNPs, tLNPs can contain lipid-conjugated antibody species whose amphiphilic character may present a greater solubilization challenge. Because the selected conditions provided robust chromatographic performance without evidence of analyte loss or column deterioration, they were adopted as a broadly applicable platform condition for subsequent method evaluation. Throughout method development, the same column was subjected to approximately 500 tLNP injections without appreciable retention time drift, peak shape deterioration, loss of analyte response, or pressure increase, supporting acceptable operational robustness under the evaluated conditions. Based on these findings, the SE-HPLC conditions were advanced to evaluation of method specificity, linearity, precision, and accuracy.

### Specificity and Linearity of RNA, Conjugated Antibody, and Free Antibody in LNP Matrix

Specificity was evaluated with empty LNPs (lacking RNA and antibody), RNA-LNPs, and antibody-functionalized RNA-LNPs (tLNPs) (**Figure 1B**). Empty LNPs produced no UV signal in the RNA or antibody quantification regions. RNA-LNPs showed a major peak at 8.3 min that matched an unencapsulated RNA standard, whereas tLNPs exhibited an additional peak at 12.5 min that was absent from the RNA-LNP control. RNA consistently exhibited a dominant peak accompanied by a smaller preceding peak. Comparable peak profiles and retention times were observed for purified RNA standards and RNA derived from tLNP samples, indicating that the feature was intrinsic to the RNA analyte rather than a consequence of nanoparticle formulation or deformulation. To facilitate antibody peak assignment, the conjugation reaction was examined in a simplified system that contained only the antibody and the conjugation reagent. The resulting reaction mixture was analyzed together with a free antibody standard and purified tLNP sample. Two distinct UV280 peaks were observed in the simplified system, matching the retention times of the conjugated-antibody peak from purified tLNPs and the free-antibody standard, respectively (**Supplemental Figure S2**). Although the difference in molecular weight between the species is only ∼2%, the conjugated antibody eluted at 12.5 min, 0.4 min earlier than the free antibody at 12.9 min (**Figure 1C**). Triplicate injections were performed to determine mean retention times and assess repeatability, with retention-time %RSD values below 0.1% for all species (**Supplementary Table S1**). The retention-time difference between conjugated and free antibody was highly reproducible across replicate injections. Together, these results demonstrate that RNA, conjugated antibody, and free antibody are chromatographically distinguishable under deformulating conditions, enabling their independent identification and quantification within tLNP formulations.

A representative RNA chromatogram is shown in **Figure 2A**. RNA response was linear from 5-200 μg/mL with R² > 0.99 and injection precision below 5% RSD at all calibration levels (**Figure 2B-C**). Back-calculated recoveries of the linear fit ranged from 99-103%, supporting quantitative analysis across the tested range (**Supplementary Table S2**). Representative normalized chromatograms at 5 μg/mL free antibody are shown in **Figure 3A**. The free antibody peak was readily detected by both UV and fluorescence, although the fluorescence trace exhibited a higher signal-to-noise ratio and a more prominent antibody peak response relative to UV280. Free antibody measurement exhibited linear response with R² > 0.99 by both UV and fluorescence detection (**Figure 3C-D).** Precision remained below 5% RSD across all calibration levels (**Figure 3B**), while fluorescence detection extended the measurable range to 1.25 μg/mL. Back-calculated recoveries of the linear fit ranged from 95-102% by UV and 90-104% by fluorescence (**Supplementary Tables S3 and S4**). Although fluorescence yielded slightly lower recovery values, its spans a wider dynamic range than UV detection. When evaluated over the same range, recoveries obtained by the two detectors were consistent. Fluorescence enabled quantification at lower antibody concentrations, but both UV and fluorescence demonstrated acceptable linearity and precision across their evaluated concentration ranges.

**Figure 2.**
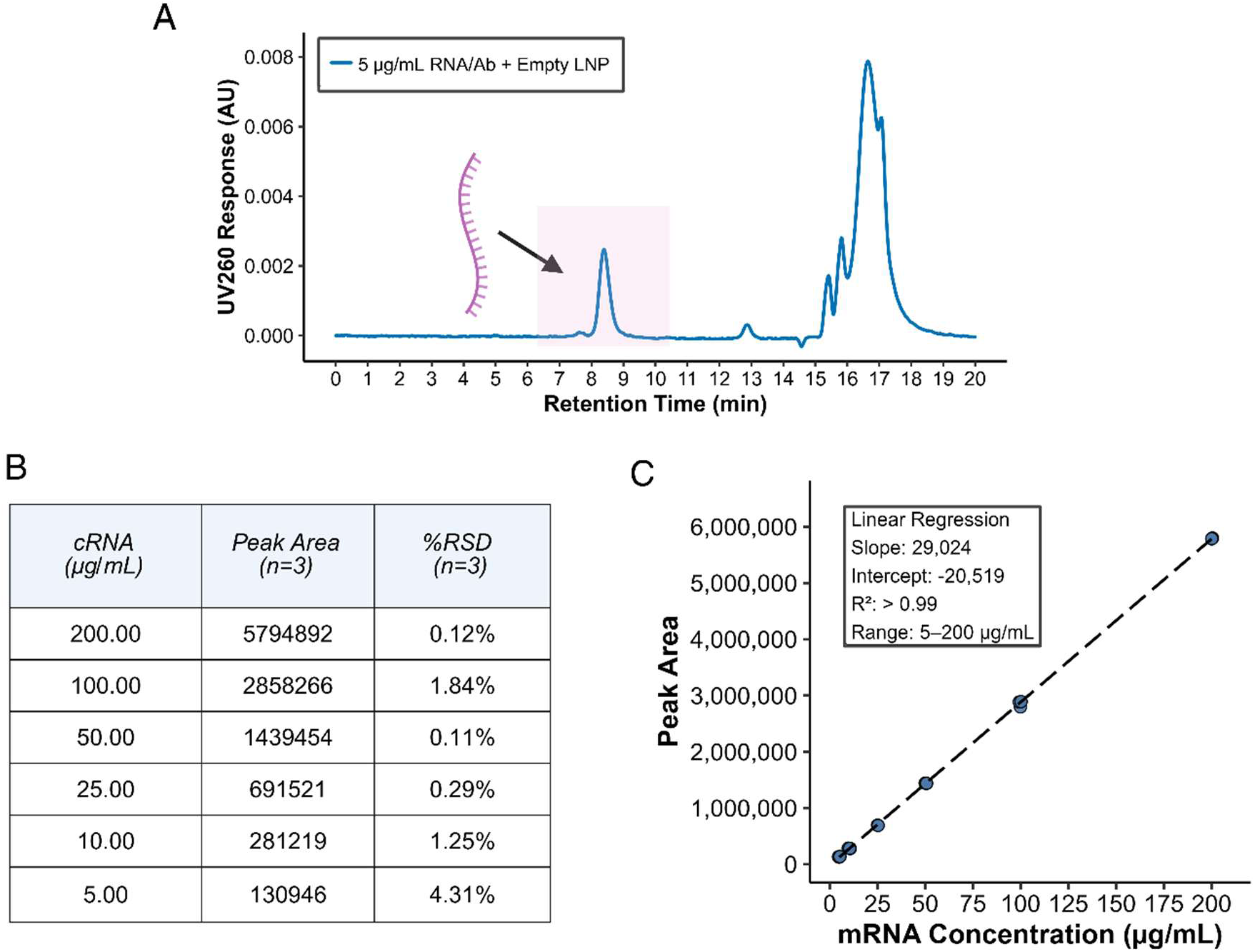
RNA calibration and quantification performance. (A) Representative UV260 chromatogram of an RNA/antibody reference standard, highlighting the RNA peak used for quantification. (B) Precision assessment for RNA quantification across the calibration range showing mean peak area and percent relative standard deviation (%RSD) calculated from triplicate injections (n = 3). (C) RNA calibration curve generated using UV260 detection across a concentration range of 5–200 µg/mL. Dashed lines represent linear regression fits used for quantitative analysis.

**Figure 3.**
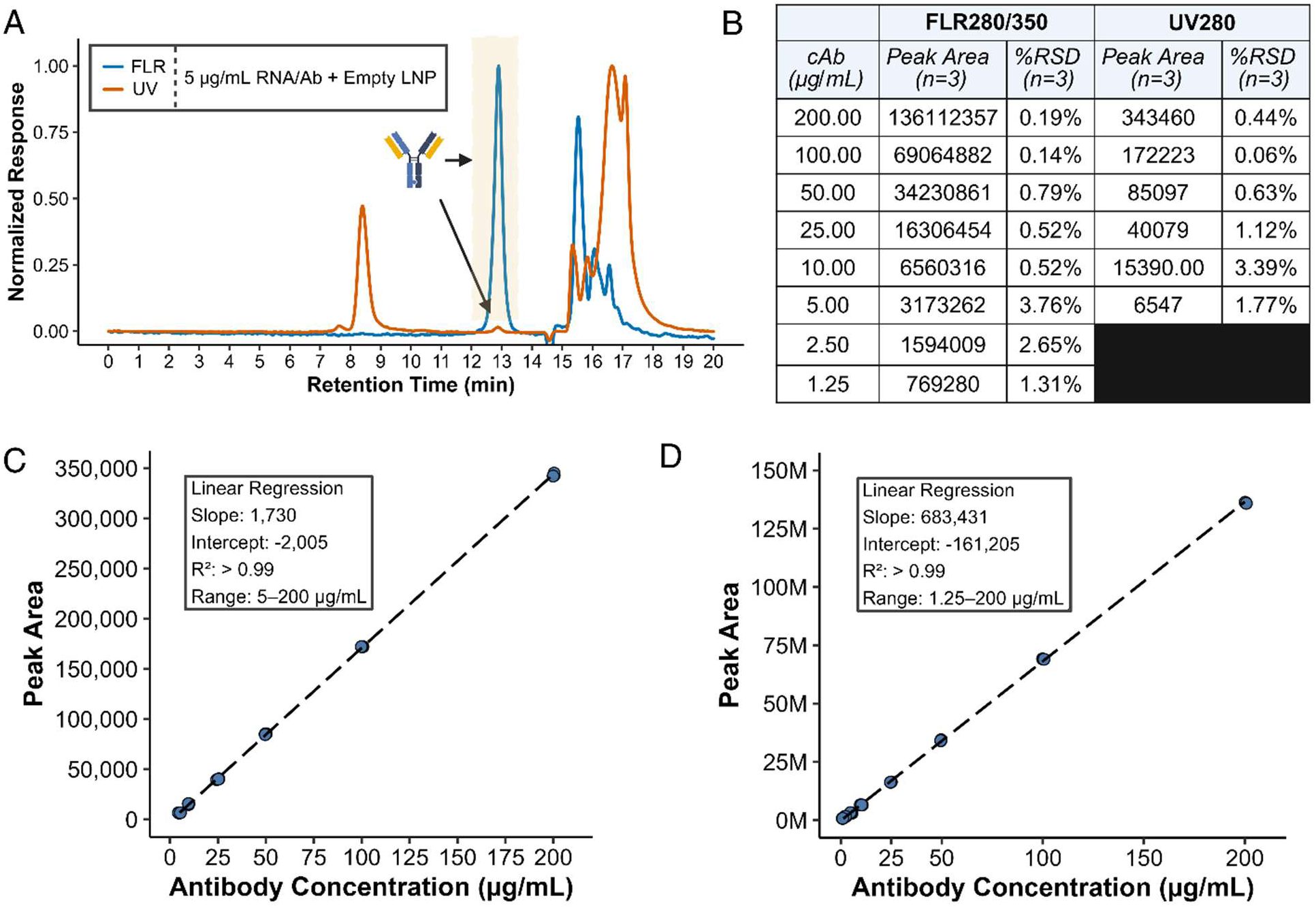
Antibody calibration and quantification performance. (A) Overlay of antibody-containing reference standards analyzed by UV280 and fluorescence (FLR280/350) detection with normalization to the highest peak in each chromatogram. The highlighted chromatographic region corresponds to the antibody peak used for quantification. Orange trace is UV. Blue trace is FLR. (B) Precision assessment for antibody quantification across the calibration range using fluorescence and UV detection. Peak area means and percent relative standard deviation (%RSD) were calculated from triplicate injections (n = 3). (C) UV280 calibration curve for antibody quantification over the range of 5–200 µg/mL. (D) Fluorescence calibration curve for antibody quantification over the range of 1.25–200 µg/mL. Dashed lines indicate linear regression fits. Calibration statistics are summarized within each panel.

### Quantification of RNA, Conjugated Antibody, and Free Antibody in tLNPs

Accuracy was assessed by spiking known amounts of RNA and free antibody into a purified tLNP sample. Calibration curves from linearity studies were applied directly to spiked and unspiked samples, with baseline RNA and antibody concentrations determined from the unspiked control. Representative chromatograms containing low and high spike levels are shown in **Figure 4A-B**. Increased RNA and antibody spike concentrations produced the expected increases in peak response without appreciable changes in peak shape or retention time. Final concentrations ranged from 64.0-113.0 μg/mL for RNA and 10.0-58.7 μg/mL for total antibody, corresponding to 12-85% free antibody (**Supplementary Table S5**). Across the evaluated range, RNA recoveries were 88-99%, while antibody recoveries were 99-112% by UV and 99-122% by fluorescence. While complete baseline separation was not achieved, the observed retention-time difference between free antibody and conjugated antibody was sufficient for quantification using perpendicular-drop integration.^28^ Triplicate injection precision remained below 5% RSD for all measurements (**Figure 4C**). UV and fluorescence detection exhibited similar overall quantitative performance. While total antibody recoveries were near 100% by both approaches, fluorescence measurements tended to yield slightly higher free antibody recoveries, particularly at lower spike concentrations, while providing total antibody recoveries closer to expected values. Collectively, these results demonstrate accurate and precise quantification of RNA, conjugated antibody, and free antibody in a representative tLNP matrix.

**Figure 4.**
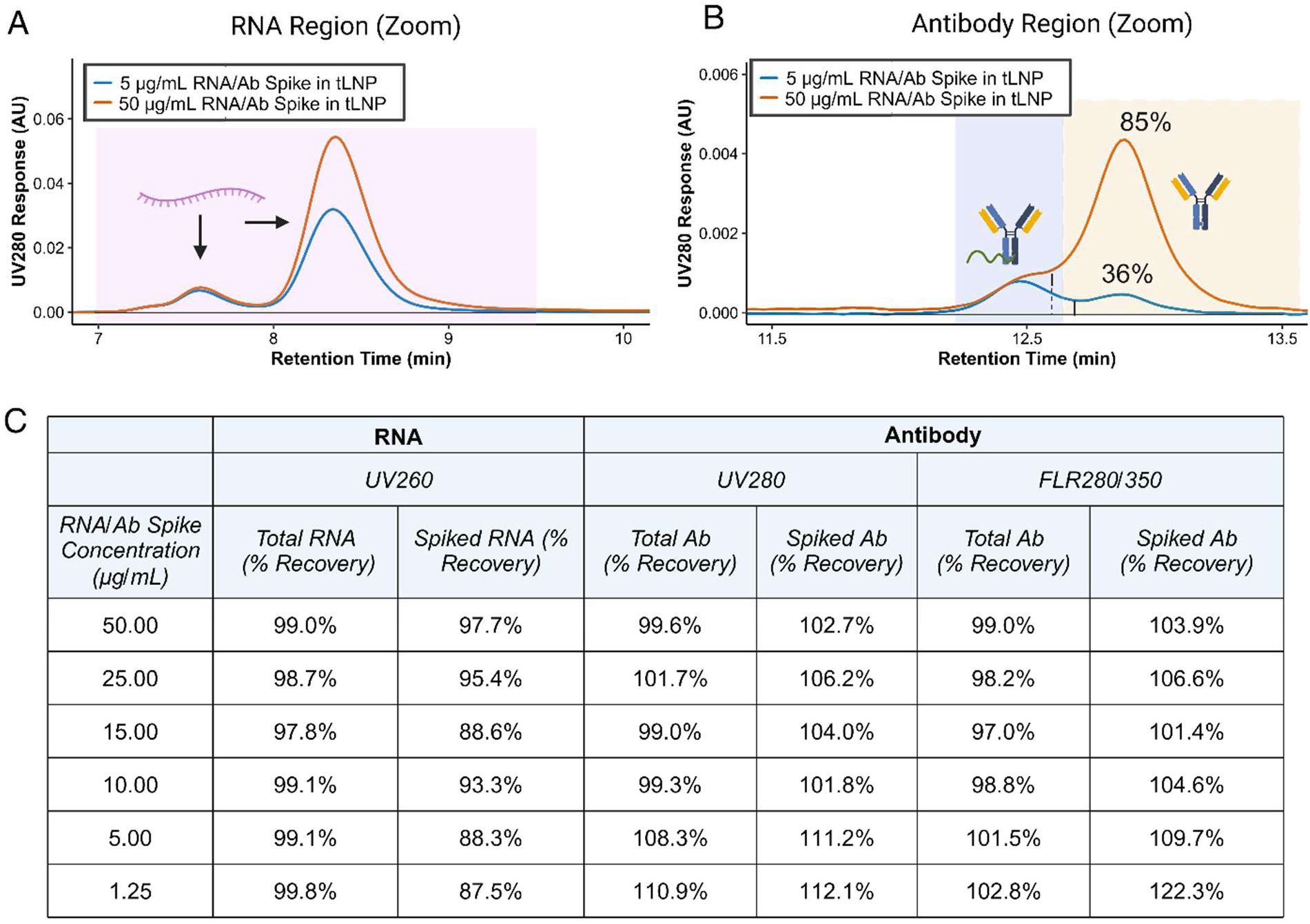
Spike-recovery evaluation of RNA and antibody quantification in antibody-conjugated lipid nanoparticles. (A) UV260 chromatograms of tLNP samples spiked with increasing concentrations of RNA and free antibody. The highlighted region corresponds to the RNA peak used for quantitative recovery assessment. (B) UV280 chromatograms of tLNP samples spiked with RNA and free antibody. Shaded regions indicate conjugated antibody and free antibody chromatographic domains used for recovery calculations. The black solid/dashed lines represent where peaks were divided for integration by perpendicular drop. (C) Percent recovery results for total and spiked RNA as well as total and spiked antibody quantified by UV280 and fluorescence (FLR280/350) detection across the evaluated spike concentration range.

With acceptable results achieved in spike-recovery experiments, we sought to challenge the method using three purified tLNP formulations. While the RNA concentration was similar, the antibody concentration was varied. The RNA and conjugated antibody peaks eluted at their expected retention times across tested formulations, indicating that increasing antibody peak area did not measurably affect chromatographic performance (**Figure 5A-B**). Importantly, no free antibody peak or shoulder was detected in any sample, providing evidence that unconjugated antibody was removed during purification. The RNA-to-antibody mass ratio provided a metric for comparing relative antibody density, with lower ratios corresponding to more conjugated antibody (**Figure 5C**). The %RSD of the tLNP samples were consistently below 2% for both RNA and conjugated antibody, indicating excellent method precision for both analytes. Together, these findings demonstrate that the SE-HPLC method can simultaneously assess RNA content, conjugated antibody content, and free antibody content in representative tLNP samples.

**Figure 5.**
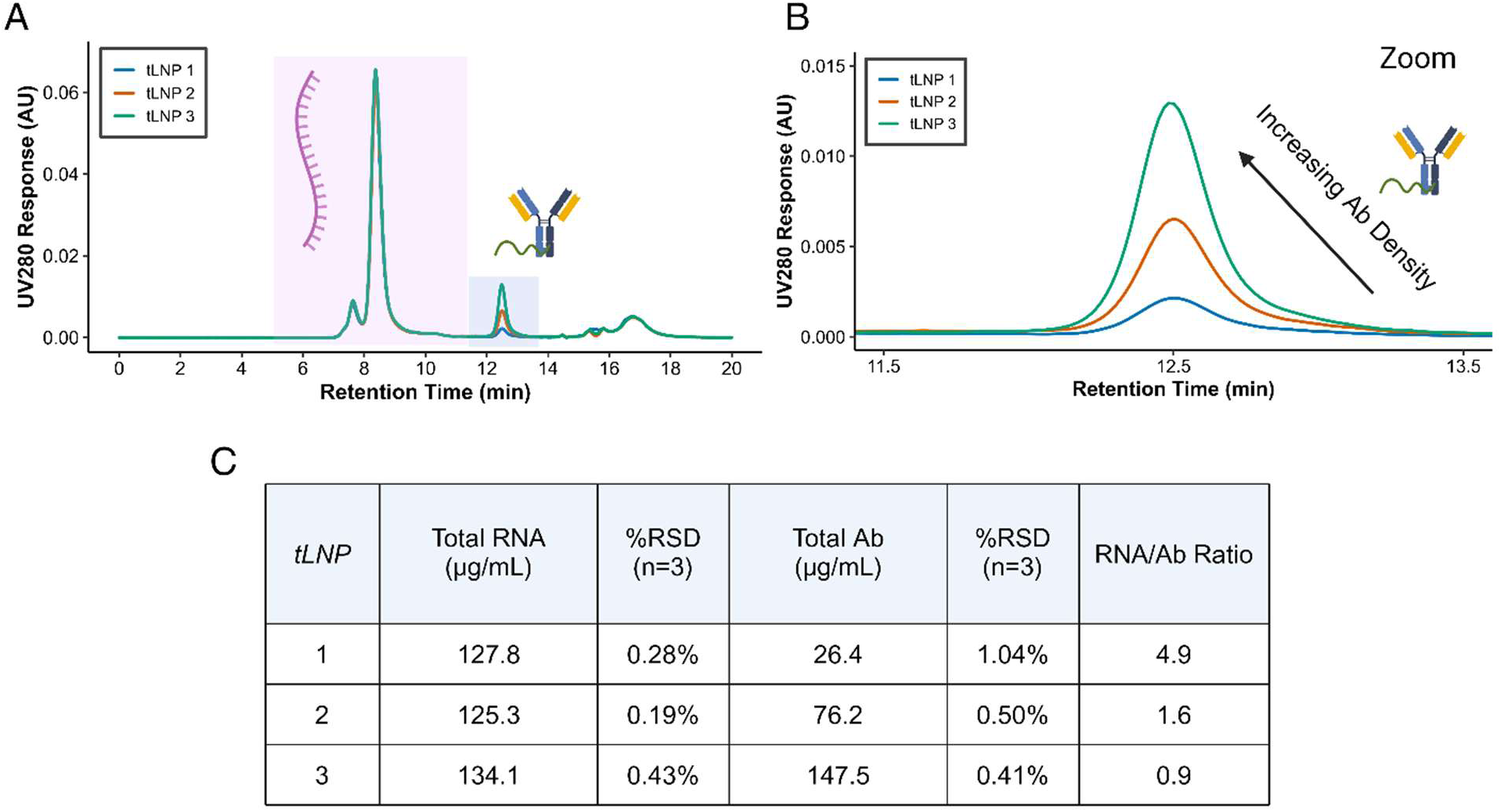
Application of deformulating SE-HPLC to antibody-conjugated lipid nanoparticles with varying antibody density. (A) UV280 chromatographic overlays of three antibody-conjugated LNP formulations containing comparable RNA content but increasing levels of surface-associated antibody. Highlighted regions correspond to the RNA and conjugated antibody peaks. (B) Expanded view of the conjugated antibody complex region demonstrating a concentration-dependent increase in antibody-associated signal with increasing antibody density. (C) Quantitative analysis of RNA and antibody content in the three tLNP formulations. Total RNA concentration, total antibody concentration, percent relative standard deviation (%RSD), and RNA:antibody ratios are reported from triplicate injections (n = 3).

## Conclusions

This work demonstrates deformulating SE-HPLC as a multi-attribute analytical method for characterization of tLNPs. By combining deformulation, chromatographic separation, and UV/fluorescence detection within a single workflow, the method enables simultaneous quantification of total RNA, conjugated antibody, and free antibody from a single injection. Furthermore, because deformulation occurs online following injection, the method minimizes pre-analytical sample handling and maintains the formulation in its native state throughout chromatographic analysis. A 5 μL injection volume was sufficient for analysis, demonstrating the method’s low sample consumption. Application to purified tLNP formulations further demonstrated the ability to discriminate between formulations with varying antibody densities, reflected by distinct RNA-to-antibody ratios. Within the concentration ranges evaluated, the method quantitatively recovered RNA and free/conjugated antibody species, supporting its utility for assessment of process yield and purification performance. While the absence of a detectable free-antibody peak in the purified tLNP samples was consistent with effective removal of free antibody, the quantitative operating range for free antibody assessment will depend on the specific targeting ligand, conjugation strategy, and analytical conditions employed. While the method was demonstrated using a full-length antibody and RNA, the general workflow may be adaptable to other targeted nanomedicine platforms with appropriate optimization of chromatographic conditions, deformulation parameters, and detection strategies. Potential applications include alternative nucleic acid payloads such as DNA, as well as targeted systems employing antibody fragments or nanobodies. Together, these findings establish deformulating SE-HPLC as a multi-attribute method for simultaneous assessment of RNA, conjugated antibody, and free antibody in antibody-functionalized LNPs.

## Author Contributions

J.R.F. developed the SE-HPLC method, performed experiments, analyzed data, prepared figures, and drafted the manuscript. A.M., G.R., and H.Z. contributed to experimental design, data interpretation, and critical review of the manuscript. P.G. and M.W. formulated LNP batches and contributed to data interpretation. H.D., and L.X. contributed to data interpretation. P.B. contributed to data interpretation and provided critical review of the manuscript. All authors reviewed and approved the final version of the manuscript.

## Acknowledgements

The authors gratefully acknowledge early development efforts at ModeX Therapeutics to advance our tLNP platform and enable the studies herein. The authors thank the ModeX Therapeutics leadership team for their support and commitment to analytical innovation. Figures were created using BioRender.com and PyMOL.

## Supplemental Figures

**Figure S1.**
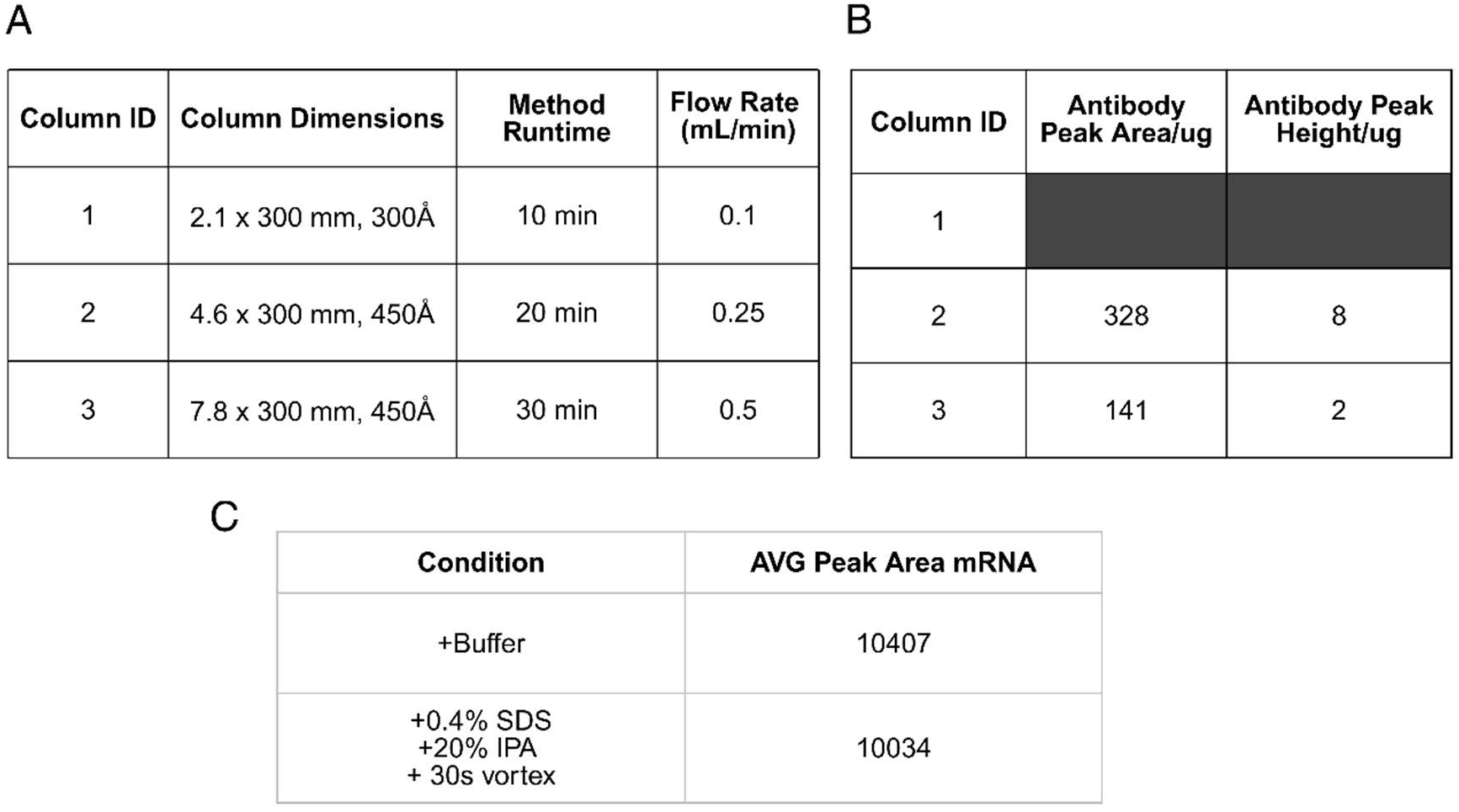
Column scouting and suitability assessment for online deformulation and quantification. (A) Three SEC column formats were evaluated to identify a configuration capable of resolving RNA and antibody species following online deformulation. (B) Relative sensitivity of each column configuration expressed as normalized peak area and peak height per unit mass injected. *ND collected for Column 1, as it did not provide sufficient resolution between RNA and antibody peaks, whereas Columns 2 and 3 provided acceptable separation for quantitative analysis. (C) Comparison of online deformulation and conventional offline deformulation using SDS/IPA followed by vortex mixing. Comparable RNA peak areas were obtained for both approaches, demonstrating that online deformulation produces equivalent analytical recovery while simplifying sample preparation.

**Figure S2.**
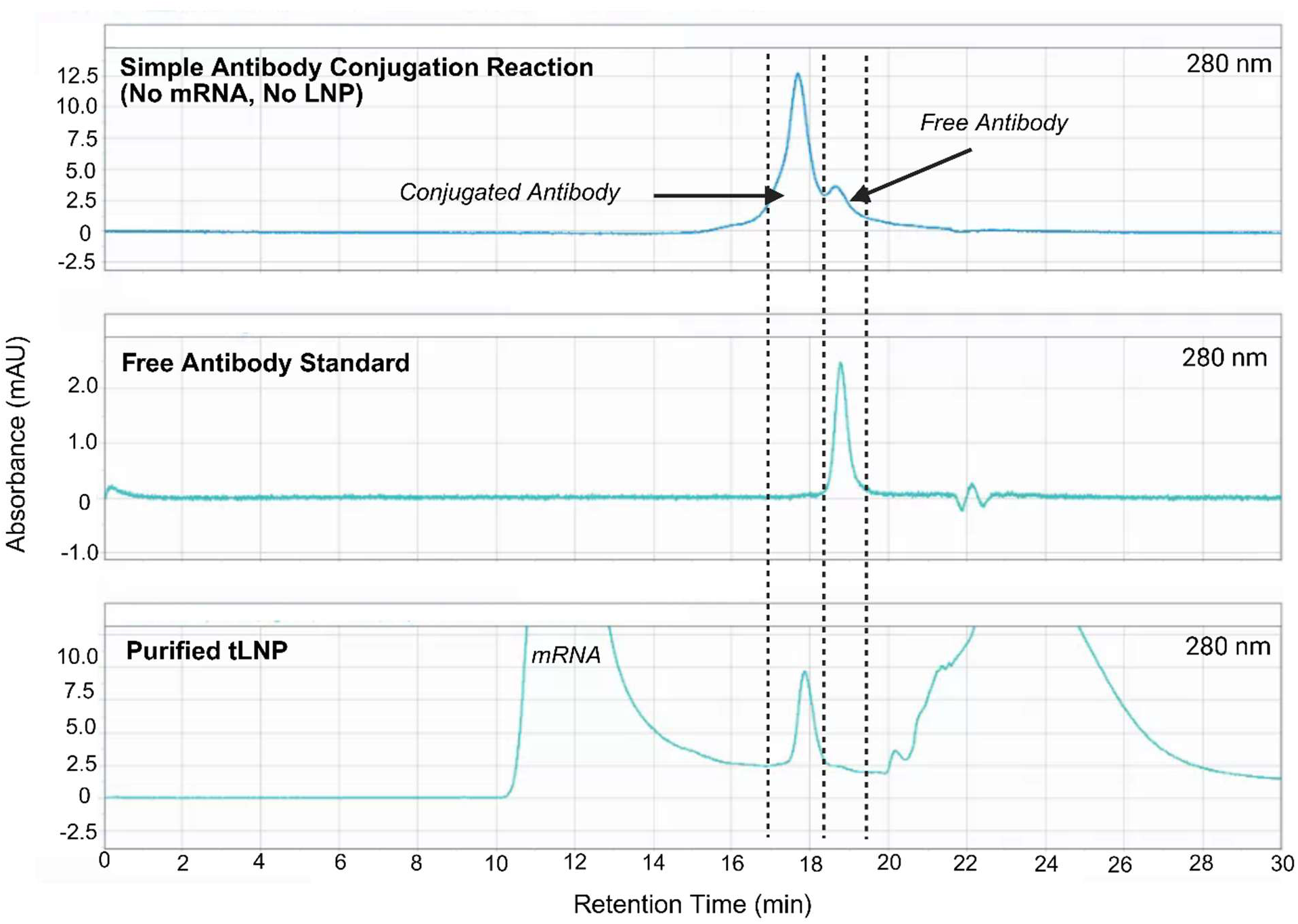
Chromatographic assignment of lipid-associated antibody species. The antibody conjugation reaction (top) yielded two distinct peaks corresponding to conjugated and unconjugated antibody forms. A free antibody standard (middle) exhibited a retention time corresponding to the later-eluting species. A representative antibody-conjugated lipid nanoparticle (tLNP) sample (bottom) exhibited an antibody-associated peak coeluting with the conjugated antibody conjugate standard, supporting assignment of the conjugated antibody complex peak observed in deformulated tLNP samples.

**Table S1.**
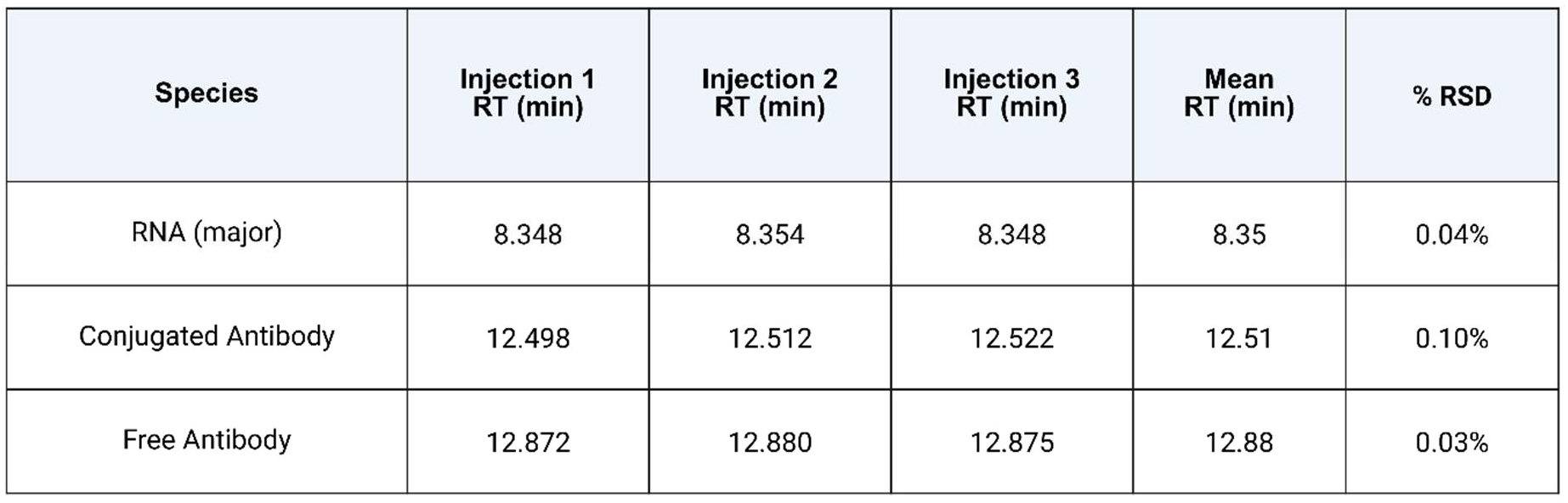
Retention time repeatability for the major RNA peak, conjugated antibody, and free antibody. Mean retention times and percent relative standard deviations (%RSD) were calculated from triplicate injections (n = 3). RNA eluted at a mean retention time of 8.35 min. Free and conjugated antibody species eluted at mean retention times of 12.88 and 12.51 min, respectively, corresponding to a retention time difference of approximately 0.4 min. The low %RSD values observed for all species indicate excellent retention time reproducibility.

| **Species** | **Injection 1<br>RT (min)** | **Injection 2<br>RT (min)** | **Injection 3<br>RT (min)** | **Mean<br>RT (min)** | **% RSD** |
|---|---|---|---|---|---|
| RNA (major) | 8.348 | 8.354 | 8.348 | 8.35 | 0.04% |
| Conjugated Antibody | 12.498 | 12.512 | 12.522 | 12.51 | 0.10% |
| Free Antibody | 12.872 | 12.880 | 12.875 | 12.88 | 0.03% |

**Table S2.** Full injection precision for RNA quantification using UV detection. Peak area data obtained from triplicate injections (n = 3) across the calibration range are summarized. Mean peak areas, standard deviations (SD), and percent relative standard deviations (%RSD) are reported for RNA quantified by UV260 detection. Low %RSD values across the calibration range demonstrate acceptable injection precision for both assays. The back calculated recovery (%) of the linear fit shows between 95%-103% across both RNA and antibody.

| **RNA - UV260** |  |  |  |  |  |  |  |
|---|---|---|---|---|---|---|---|
| *cRNA<br>(ug/mL)* | *Injection 1* | *Injection 2* | *Injection 3* | *Mean Peak Area* | *SD* | %RSD | %Back Calculated Recovery |
| 200.00 | 5795918 | 5801567 | 5787192 | 5794892 | 7242 | 0.12% | 100.17 |
| 100.00 | 2797565 | 2886070 | 2891164 | 2858266 | 52630 | 1.84% | 99.31 |
| 50.00 | 1441319 | 1438846 | 1438199 | 1439454 | 1646 | 0.11% | 99.31 |
| 25.00 | 693333 | 691886 | 689344 | 691521 | 2019 | 0.29% | 102.93 |
| 10.00 | 277659 | 284674 | 281325 | 281219 | 3508 | 1.25% | 99.40 |
| 5.00 | 124628 | 132718 | 135492 | 130946 | 5644 | 4.31% | 101.52 |

**Table S3.** Full injection precision for antibody quantification using UV detection. Peak area data obtained from triplicate injections (n = 3) across the calibration range are summarized. Mean peak areas, standard deviations (SD), and percent relative standard deviations (%RSD) are reported for antibody quantified by UV280 detection. Low %RSD values across the calibration range demonstrate acceptable injection precision for both assays. The back calculated recovery (%) of the linear fit shows between 95%-103% across both RNA and antibody.

| **Antibody - UV280** |  |  |  |  |  |  |  |
|---|---|---|---|---|---|---|---|
| *cAb (ug/mL)* | *Injection 1* | *Injection 2* | *Injection 3* | *Mean Peak Area* | *SD* | %RSD | %Back Calculated Recovery |
| 200.00 | 342918 | 345179 | 342283 | 343460 | 1522 | 0.44% | 99.82 |
| 100.00 | 172163 | 172334 | 172172 | 172223 | 96 | 0.06% | 100.83 |
| 50.00 | 85460 | 85352 | 84479 | 85097 | 537 | 0.63% | 99.39 |
| 25.00 | 39593 | 40474 | 40170 | 40079 | 447 | 1.12% | 101.96 |
| 10.00 | 14788 | 15663 | 15719 | 15390 | 522 | 3.39% | 95.91 |
| 5.00 | 6414 | 6601 | 6627 | 6547 | 116 | 1.77% | 95.16 |

**Table S4.** Full injection precision for antibody quantification using fluorescence detection. Peak areas obtained from triplicate injections (n = 3) across the calibration range are shown together with the calculated mean peak area, standard deviation (SD), and percent relative standard deviation (%RSD). Precision remained acceptable throughout the evaluated concentration range, including low-concentration standards used to assess method sensitivity. The calculated back recovery of the linear fit is between 90-104% across a wide dynamic range.

| **Antibody - FLR280/350** |  |  |  |  |  |  |  |
|---|---|---|---|---|---|---|---|
| *cAb<br>(ug/mL)* | *Injection 1* | *Injection 2* | *Injection 3* | *Mean Peak Area* | *SD* | %RSD | %Back Calculated Recovery |
| 200.00 | 136149071 | 136347530 | 135840472 | 136112357 | 255514 | 0.19% | 100.30 |
| 100.00 | 69178818 | 69014645 | 69001184 | 69064882 | 98900 | 0.14% | 98.72 |
| 50.00 | 34533719 | 34014040 | 34144824 | 34230861 | 270311 | 0.79% | 99.36 |
| 25.00 | 16233725 | 16400243 | 16285395 | 16306454 | 85233 | 0.52% | 103.79 |
| 10.00 | 6522729 | 6589221 | 6569000 | 6560316 | 34085 | 0.52% | 101.72 |
| 5.00 | 3042987 | 3199706 | 3277094 | 3173262 | 119272 | 3.76% | 102.61 |
| 2.50 | 1545411 | 1621250 | 1615367 | 1594009 | 42190 | 2.65% | 97.07 |
| 1.25 | 764257 | 762724 | 780859 | 769280 | 10056 | 1.31% | 90.10 |

**Table S5.** Known concentrations for varying spike levels and resulting percentage of free antibody. The total RNA concentration varied between 64.3-113 ug/mL in the tested spike levels. The total antibody concentration varied between 10.0-58.7 ug/mL, with the percentage of free antibody ranging from 12.5% to 85.1%, calculated from the expected mass of each antibody species at the spike level.

| **Total RNA Concentration (ug/mL)** | **Total Antibody Concentration (ug/mL)** | **Conj Antibody Concentration (ug/mL)** | **RNA/Antibody Spike Final Concentration (ug/mL)** | **% Free Ab** |
|---|---|---|---|---|
| 113 | 58.7 | 8.735 | 50 | 85.1 |
| 88 | 33.7 | 8.735 | 25 | 74.1 |
| 78 | 23.7 | 8.735 | 15 | 63.2 |
| 73 | 18.7 | 8.735 | 10 | 53.4 |
| 68.0 | 13.7 | 8.735 | 5 | 36.4 |
| 64.3 | 10.0 | 8.735 | 1.25 | 12.5 |

## Notes

### Competing Interest Statement

Authors are employees of ModeX Therapeutics.

